# Two Liberibacter Effectors Combine to Suppress Critical Innate Immune Defenses and Facilitate Huanglongbing Pathogenesis in Citrus

**DOI:** 10.1101/2021.11.26.470170

**Authors:** Supratim Basu, Loan Huynh, Shujian Zhang, Roel Rabara, Hau Nguyen, Jeanette Valesquez, Guixia Hao, Godfrey Miles, Qingchun Shi, Ed Stover, Goutam Gupta

## Abstract

Genome sequence analyses predicted the presence of effectors in the gram-negative *Candidatus* Liberibacter asiaticus (*C*Las) even without the presence of a classical type III secretion system. Since *CLas* is not culturable, it is not possible to perform traditional gene knockout experiments to determine the role of various effectors in Huanglongbing (HLB) pathogenesis. Therefore, we followed an alternative functional genomics approach to examine the role of the *C*Las effectors in HLB pathogenesis in general and more specifically in suppressing citrus innate immune response. Here, we focused on the *C*Las effectors, P235 and Effector 3, to perform the following studies. *First*, proteomic studies by LC-MS/MS were conducted to screen the putative interacting citrus protein partners of P235 and Effector 3 from the healthy and *C*Las-infected Hamlin extracts and the most probable candidates were identified based upon their high protein scores from LC-MS/MS. *Second*, a transgenic tobacco split GFP system was designed for *in planta* detection of the most probable citrus interacting protein partners of P235 and Effector 3. *Third*, *in vitro* and *in planta* studies were performed to show that each of two effectors interacts with and inhibits the functions of multiple citrus proteins belonging to the innate immune pathways. These inhibitory interactions led to a high level of reactive oxygen species (ROS), blocking of bactericidal lipid binding protein (LTP), and induction of premature programmed cell death (PCD), thereby supporting *C*Las infection and HLB pathogenesis. *Finally*, an LTP mimic was designed to sequester and block the *C*Las effector and to rescue the bactericidal activity of LTP.

## Introduction

Huanglongbing (HLB) is the most devastating citrus disease ^1,2^. While endemic in Asia for over a century ^3,4^, HLB was first encountered about a 15 years ago in Florida with the emergence of Asian Citrus Psyllid (ACP) vectors carrying HLB-causing *Candidatus* Liberibacter asiaticus (*CLas*). Since then, HLB has been widespread in Florida and is looming large on California and Texas, the two other citrus producing states in the US. Robust HLB management tools are urgently needed for sustaining a productive and profitable citrus industry^5^. These tools include development of both bactericidal and anti-infective molecules for HLB treatment. Recently, we reported the development of novel citrus-derived *CLas*-killer peptides that can be used for HLB treatment by topical delivery ^6^. In this study, we focused first on identifying the critical steps associated with the breakdown of citrus innate immune defense in response by the *CLas* effectors and then on developing therapeutic and anti-infective molecules to block them. Typically, the plant innate immune defense involves multiple pathways including pathogen or microbe-associated molecular pattern (PAMP or MAMP) triggered immunity (PTI or MTI), effector triggered immunity (ETI), and plant hormone, such as salicylic (SA), jasmonic acid (JA), and ethylene (ET) induced immunity^7–12^. The PTI or MTI provides the first line of plant defense against pathogens or microbes through the recognition of PAMP or MAMP, such as bacterial liposaccharide (LPS), elongation factor thermal unstable (EF-Tu), flagellin. PAMP or MAP recognition is mediated by the plasma membrane pattern recognition receptors (PRR) that include leucine-rich receptors (LRR), flagellin receptor (FLS2), EF-Tu receptor (EFR). The plasma membrane PRR recognition induces intracellular mitogen-associated protein kinase (MAPK) signaling leading to the expression of pathogen-related (PR) or defense genes ^13–15^. However, pathogen effectors can block both intracellular and extracellular steps in the PTI pathway ^16,17^. To counter pathogen induced blocking of the PTI pathway, plants have evolved the ETI pathway in which intracellular nod-like receptors (NLR) recognize the pathogen effectors and augment the MAPK signaling and PR gene expression. The ETI pathway also induces hypersensitive response through the production of reactive oxygen species (ROS), which causes cell death at site of infection thereby limiting pathogen spread. The PTI and ETI pathways also couple to intracellular plant hormone SA/JA/ET pathways, which also involve ROS production and induction of PR genes. It has been demonstrated that the effectors from plant pathogenic bacteria can inhibit one or more steps in these pathways ^18–21^. Also, the bacterial effectors are known to subvert multiple steps leading to programmed cell death (PCD) in plant, which is a form of immune defense by PTI and/or ETI to control infection ^22–24^. Therefore, it was of interest for us to determine which steps in the citrus innate immune defense are affected by the *CLas* effectors. Note that the *CLas* effectors are smaller in number because of the small (1Mb) genome-size ^25,26^ and are also unique in that the bacterium does not have a type III or VI secretion system like many other plant pathogenic gram-negative bacteria ^27,28^. Gram-negative bacteria with 5 Mb genomes have several hundred unique effectors ^29^ as opposed to only 20 effectors identified, so far, from *CLas* ^30,31^. However, interactome studies revealed that even an effector from a gram-negative bacterium carrying hundreds of effectors can target more than one protein from the host plant ^17,32^. Therefore, it was of interest to examine whether each *CLas* effector may target multiple citrus proteins to effectively suppress the innate immune defense and establish infection.

In this study, we focused on two *CLas* effectors, P235 and Effector 3. *First*, we performed *in vitro* and *in planta* studies to identify the prominent citrus proteins targeted by P235 and Effector 3. *Second*, we performed functional assays to determine whether P235 and Effector 3 have inhibitory effects on their citrus protein targets. Next, we performed molecular dynamic simulations to analyze the details of interaction between P235 (and Effector 3) and their selected citrus targets and predicted which pairwise interactions are critical for inhibition of the citrus target function. *Finally*, we validated our prediction of the inhibitory mechanism by site-specific mutations on the citrus protein(s) that affect the critical pairwise interactions. We discovered that each of the two effectors can directly target several citrus proteins, which belong to the innate immune networks. A clear understanding of the inhibitory mechanisms will provide guidelines for countering *CLas* effectors and developing anti-infectives to block HLB pathogenesis.

## Methods

### Experimental Procedures

#### Plant Materials and growth conditions

Hamlin trees verified as being HLB-free and ACP-friendly were purchased and placed in the green house. One branch cage placed in the upper part of each tree (3 replicates) was filled with 75ACP from an infected population while other trees had cages with clean 75ACPs placed serving as control. The insects were allowed to feed on the trees for a week and then the insects were killed by spraying with topical insecticide. The ACPs were tested for *C.Las* and the trees were subsequently returned to the greenhouse. Leaf samples were collected from the untreated and infected plants and flash frozen in liquid nitrogen and stored for further analysis.

#### Cloning and overexpression of effectors and targets in *E. coli*

The effectors from *Liberibacter asiaticus* were identified, codon optimized and cloned in pUC57 by GenScript. The effectors were then amplified and cloned in pET28(a) vector between NdeI and BamHI sites. The positive clones were inoculated overnight in LB with Kanamycin . The overnight culture (1%) was grown until the OD reached 0.6 and then induced with IPTG at 30°C for Overnight. Cells were harvested next day and resuspended in protein isolation buffer (20mM Tris-Cl,7.4, 150mM NaCl and 10% glycerol). The cell suspension was sonicated and centrifuged at 14000 rpm, 4°C, 30 mins. Supernatant was collected and the inclusion bodies were treated with 9M urea. Following treatment with urea the cell suspension was centrifuged and supernatant was collected and refolded. The refolded protein from the inclusion bodies and the soluble fractions were purified using TALON metal affinity resin.

#### Isolation of total protein from citrus

Fresh leaf tissue, from five Hamlin trees (Citrus sinensis L. Osbeck) was pulverized in liquid nitrogen using a pestle and mortar and the resulting fine powder stirred with 1.5 volumes of extraction buffer (50 mM HEPES pH 7.5, 5 mM EDTA, 5 mM EGTA, 10 mM dithiothreitol (DTT), 10% glycerol, 7.5% polyvinylpolypyrolidone (PVPP), and a protease inhibitor cocktail, Complete™, Boehringer Mannheim). The slurry was subsequently mixed on a reciprocating shaker (100 oscillations per min) for 10 min, at 4°C, followed by centrifugation 15,000 g for 30 min at 4°C. The supernatant was removed and immediately flash-frozen in liquid nitrogen and stored at -80°C until needed.

#### Pull down assay and LC-MS/MS analysis to identify citrus targets

The purified refolded effector proteins were incubated with total protein (15μg) isolated from healthy and infected citrus leaf extract for 2 h at 4°C. The effector-protein complex was incubated with TALON metal affinity resin at 4°C overnight. The resin was washed with column buffer (50mM Tris-Cl, pH7.4, 150mM, 10% glycerol) and eluted with imidazole (250mM). The eluted protein complex was sent for LC-MS/MS analysis to identify the citrus targets. The spectra were searched against the Uniprot database, and taxonomy was set to *Citrus sinensis*. Only peptides that were ranked 1 were selected and finally those targets were selected for further analysis that had a 95% confidence ^33^.

### Enzymatic Assays and their Inhibitions by the CLas effectors

#### Superoxide dismutase assay

Superoxide dismutase assay was quantified based on its ability to inhibit the photochemical reduction of nitroblue tetrazolium (NBT) by superoxide radical and assayed following (Superoxide Dismutase Kit; Catalog Number: 7500-100-K) with some modifications. The reaction mixture (3ml) contained 13 mM methionine, 75 mM NBT, 2 mM riboflavin,100 mM EDTA and 0.3ml leaf extracts. The volume was made up to 3ml using 50 mM phosphate buffer with the addition of riboflavin at the very end. One the reaction mixture is made they were mixed well and incubated below two 15-W fluorescent tubes with a photon flux density of around 40 mmol m^-2^ s^-1^ for 10 mins. Once the reaction is completed, the tubes were covered with a black cloth and the absorbance was measured at 560 nm. The non-irradiated mixture served as control and the absorbance so measured is inversely proportional to the amount of enzyme added. SOD activity is defined as the amount of enzyme that caused 50% inhibition of the enzymatic reaction in the absence of the enzyme.

#### Aspartyl protease assay

The protease assay was performed using a fluorescence based (BODiPY) EnzChek protease assay kit. The analysis of aspartyl protease activity was done by incubating it with no other proteins in sodium citrate buffer (50 mM, pH 4.5). To perform the inhibitory effect of P235 on the protease activity the renatured aspartyl protease was preincubated with increasing concentrations of P235 at 4°C for 2 h in sodium citrate buffer. Following incubation, BODiPY-labeled casein substrate was added, and the reaction was monitored by measuring fluorescence in Tecan Infinite 200 PRO microplate reader at 485±12.5 nm excitation/530±15 nm emission filter. The assays were conducted in replicates.

#### Glycosyl hydrolase Assay

The inhibitory effect of recombinant P235 on recombinant glycosyl hydrolase was assayed using the β-Glucosidase Activity Assay Kit (MAK129, Sigma). Enzymatic reactions were carried out in K-Phosphate buffer (100 mM, pH 6.5) with *p*-nitrophenyl-β-D-glucopyranoside (ß-NPG) for 20 minutes at 37 °C. Final absorbance of the hydrolyzed product was measured at 405 nm.

#### Aldehyde dehydrogenase assay

This assay was performed using ALDH Activity Abcam Assay Kit with modifications. In short, purified aldehyde dehydrogenase was incubated with increasing concentration of substrate (acetaldehyde) for 1 h. The absorbance was measured at 450nm and expressed in terms of NADH standard as mU/ml.

#### Trypsin inhibition assay

The trypsin inhibition assay was done in triplicate and the result is expressed as a means of three replicates. In short, residual trypsin activity was measured by monitoring the change in absorbance at 247 nm in presence of increasing concentration of recombinant purified Kunitz Trypsin inhibitor (KTI) when incubated with *p*-toluene-sulfonyl-L-Arg methyl ester (Sigma). To study the inhibitory action of P235 on KTI action, increasing molar concentration of P235 was mixed with BSA and KTI and incubated for 1 h at room temperature. The result was analyzed using SDS PAGE.

#### *In-planta* split GFP assay (agro-infiltration)

*Agrobacterium tumefaciens* LBA4404 transformant cells carrying effectors P235, Effector 3 and the targets from citrus plants (Aspartyl protease, glycosyl hydrolase, superoxide dismutase, Kunitz trypsin inhibitor protein, lectin etc) respectively are cloned in pR101 vector and cultured overnight in LB medium with 50 μg ml^−1^ of rifampicin and 50 μg ml^−1^ kanamycin and resuspended in 10 mM MgCl_2_, 10mM MES. The culture was diluted to an optical density of 0.5 (OD 600nm). For each effector-target interaction, three leaves of 4 *N. benthamiana* plants overexpressing GFP1-9 were infiltrated with the *A. tumefaciens* suspension containing the effector and the target plasmids respectively. The agro-infiltrated leaves were analyzed for protein localization at 3 dpi under a microscope (Olympus BX51-P) equipped with a UV light source. Agroinfiltrated plants were kept in a greenhouse for 24 h and the interaction was visualized using Illumatool lighting system (LT-9500; Lightools Research) with 488 nm excitation filter (blue) and a colored glass 520 nm long pass filter. The photographs were taken by Photometric CoolSNAP HQ camera.

#### Estimation of superoxide anion

Leaf discs from agro-infiltrated tobacco plants were incubated at 25°C on a shaker for 30 mins in dark in 1 ml of K-phosphate buffer (20 mM, pH 6.0) containing 500 µM XTT. The increase in absorbance was measured at 470nm in a spectrophotometer.

#### Lipid Binding and MIC Assays for LTP

Lipid binding activity of recombinant LTP-6X His protein overexpressed and purified from *E. coli* was mixed with of 2-*p*-toluidinonaphthalene-6-sulphonate (TNS) at 25 °C. The results were recorded at excitation 320nm and the emission at 437nm. The inhibitory action of P235 on LTP was assessed using increasing concentration of P235 and the results were measured. Purified GFP was used as a control. The minimum inhibitory concentration (MIC) of the LTP (lipid transfer protein) was performed using broth microdilution technique. The assay was carried out using 5 × 10^5^ colony forming units (CFU/ml) in MHB. MIC was defined as the lowest concentration of the protein required to inhibit the visible growth of bacterial strains used.

#### Estimation of ion leakage from leaf discs

*Agrobacterium tumefaciens* LBA4404 transformant cells carrying Effector 3 and the targets from citrus plants Kunitz trypsin inhibitor protein cloned in pR101 vector was cultured overnight in LB medium with 50 μg ml^−1^ of rifampicin and 50 μg ml^−1^ kanamycin and resuspended in 10 mM MgCl_2_, 10mM MES. The culture was diluted to an optical density of 0.5 (OD 600nm). For the assay, three leaves of N*. benthamiana* plants previously treated with paraquat (100µM) were infiltrated with the *A. tumefaciens* suspension containing the effector alone, Kunitz alone and the mixture of effector 3 and Kunitz respectively ^30,31^ and incubated for 48h. Leaf discs were prepared by punching the leaf discs with a cork puncher. The punctured leaf discs were placed in water (50 mL) for 5 minutes to mitigate the error of measuring ion leakage due to injury inflicted on the leaves due to puncturing. Following, preincubation the leaf discs were incubated in 5 mL of water for 3h. Conductivity was measured after 3h using Mi180 bench meter and this value is referred to as A. Leaf discs with the bathing solution were then incubated at 95°C for 25minutes and then cooled to room temperature to enable complete ion leakage. The conductivity was measured again, and this value is referred to as B. Ion leakage is subsequently expressed as (value A/ value B) x100. All the experiments were carried out in three biological replicates with five leaf discs for each sample^34,35^.

### Pathogen inoculation and LTP treatment in *N.benthamiana* leaves

*Pseudomonas syringae pv. Tomato* DC3000 was cultured on King’S B (KB) medium containing 50 μg mL^−1^ rifampicin. Overnight, log-phase cultures were grown to an optical density at OD_600 nm_ of 0.6 to 0.8 (OD 0.1 = 10^8^ cfu mL^−1^) and diluted with 10 mM MgCl_2_ to the concentrations of 10^5^ cfu mL^−1^ before inoculation. Control was performed with 10 mM MgCl_2_. The bacterial suspensions were infiltrated into the abaxial surface of a leaf using a 1-mL syringe without a needle. *Agrobacterium tumefaciens* LBA4404 transformant cells carrying P235 and LTP protein cloned in pR101 vector was cultured overnight in LB medium with 50 μg ml^−1^ of rifampicin and 50 μg ml^−1^ kanamycin and resuspended in 10 mM MgCl_2_, 10mM MES. The culture was diluted to an optical density of 0.5 (OD 600nm). For the assay, infected leaves of N*. benthamiana* plants were infiltrated with the *A. tumefaciens* suspension containing the LTP alone, LTP+P235 alone, different mimics ^36^.

### Molecular Modeling

#### Prediction of protein 3D structures and complexes

3D structures of the two *CLas* effector (P235 and Effector 3) and the two citrus proteins (LTP and KTI) were predicted using I-TASSER (https://zhanglab.ccmb.med.umich.edu/I-TASSER/). We then used HADDOCK version 2.2 webserver to predict interaction interfaces of P235-LTP and Kunitz-E3 complexes (http://milou.science.uu.nl/services/HADDOCK2.2/). Selected complexes of P235-LTP and Kunitz-E3 were further refined using MD simulations of these complexes in the presence of water

#### Protein-water system setup for MD simulation

Our simulations started with single protein (*i.e.* LTP, P235, Kunitz, or E3) in water. These systems contained 10,000 water in a box of 6.9 × 6.8 × 7.1 nm^3^. To refine the models of P235-LTP and Kunitz-E3 obtained from HADDOCK. We conducted MD simulations of these complexes in the presence of water. The protein-protein complex systems contain 30,000 water in a box of 9.9 × 9.9 × 9.9 nm^3^ with excess NaCl at 150 mM to mimic experimental conditions. For Kunitz-E3 complexes, we focus on model with Kunitz’s active loop in close contact with E3’s interface that contain either aspartic acid or glutamic acid residues or a large hydrophobic surface. For P235-LTP complexes, we focus on model with LTP’s lipid entrance site B1 and B2 (see Fig. S2) in close contact with P235. Following MD simulation, systems with stable complexes and adequate protein-protein pairwise residues interactions were then further validated by extended MD simulation.

#### Protein-bilayer system setup for MD simulation

Our simulations started with a single LTP in the water and a mimetic of the *E. coli* inner membrane composed of a 3:1 ratio of 1-palmitoyl-2-oleoyl-*sn*-glycero-3-phosphoethanolamine^34^ (POPE) and 1-palmitoyl-2-oleoyl-*sn*-glycero-3-phosphoglycerol (POPG). Lipid bilayers are constructed with the Charm-GUI membrane builder^37^ followed by 40 ns of NpT simulation at 310 K with semi-isotropic pressure coupling. The LTP-bilayer system contained 10,000 water molecules and 128 lipid molecules in a box of 6.1 × 6.1 × 12.5 nm. We also conducted simulation of P235-LTP complex in the bilayer POPE: POPG (3:1 ratio) to further refine the P235-LTP models obtained from MD simulation of the P235-LTP complexes in the water. The P235-LTP/bilayer contained 23,600 water molecules an’d 256 lipid molecules in a box of 8.7 × 8.7 × 13.7 nm. LTP, or P235-LTP complex was placed 3.5 nm away from the center of mass of the lipid bilayer along its normal. Protein/bilayer systems were neutralized and excess NaCl was added at 150 mM to mimic experimental conditions.

#### Simulation protocol

For MD simulations, the TIP3P water model was used. with CHARMM modifications ^38^. Water molecules were rigidified with SETTLE ^39^ . and molecular bond-lengths were constrained with P-LINC Lennard-Jones interactions ^40^ were evaluated using a group-based cutoff, truncated at 1 nm without a smoothing function. Coulomb interactions were calculated using the smooth particle-mesh Ewald method^41–43^ with a Fourier grid spacing of 0.12 nm ^44^. Simulation in the *NpT* ensemble was achieved by semi-isotropic coupling at 1 bar with coupling constants of 4 ps ^45,46^ and temperature-coupling the simulation system using velocity Langevin dynamics with a coupling constant of 1 ps ^47^. The integration time step was 2 fs. The non-bonded pair-list was updated every 20 fs ^48^.

## Results

### *In vitro* protein assay to identify the citrus protein targets of P235 and Effector 3

Effector 3 has a predicted chloroplast targeting signal sequence whereas P235 has an N-terminal nuclear localization signal (NLS) ^30,31^. Note that most of the *CLas* effector do not possess classical type III secretion signal sequence. However, they may be secreted by the type II secretion pathways probably via outer-membrane protein transporters ^49,50^. Homology modeling predicted the presence of helical bundles in the structure of P235 as shown in Fig. S1A of the supplementary material. Note that similar helical bundles are also present in AvrRps4, a *P. syringae* effector involved in plant immunity ^51^. It is suggested that the helical effectors from bacteria may interact with multiple plant helical proteins via intermolecular coiled-coil interactions ^52,53^. It was of interest to us to determine whether P235 interacted with the helical proteins from the citrus innate immune repertoire. These proteins may be located on the plasma membrane, in the cytosolic fluid or vacuole, and in the nucleus. Homology modeling also predicted 2 helix bundle in the structure of Effector 3 in addition to a disordered C-terminal segment (see Fig. S1B of the supplementary material). The latter may make Effector 3 a promiscuous binding partners of several citrus proteins. In addition, due to the presence of chloroplast targeting signal, Effector 3 may be a potential *CLas* effector. Note that multiple chloroplast proteins are involved in ROS production and plant hormone signaling ^54^, which may mediate cell death as an innate immune response. It was of interest to examine whether Effector 3 bound to any citrus chloroplast protein associated with ROS production, phytohormone signaling, or cell death. Although, we predicted a certain type of citrus target proteins for P235 and Effector 3, a whole proteome screening was needed to identify all their prominent targets.

The steps in our target identification scheme is shown in Fig. 1 (left). *First*, we expressed P235 and Effector 3 in *E. coli* with C-terminal His_6_-tags. The Effector 3 was expressed without the signal sequence. Both the proteins were extracted from the inclusion body and re-folded. *Second*, the His-tagged P235 and Effector 3 were bound to TALON columns and were incubated with citrus protein extracts from uninfected and *CLas*-infected Hamlin, which was infected by caged *CLas*-carrying psyllids. *Third*, bound citrus protein targets were eluted from the column and identified by LC-MS/MS. *Finally*, the spectra from LC-MS/MS were searched against the Uniprot database with taxonomy set to *Citrus sinensis*. The highest ranked citrus proteins, in terms of the LC-MS/MS protein score ^55^, were selected as putative targets of P235 and Effector 3. See supplementary Tables SIA and SIB for all the citrus targets of P235 and Effector 3 with high protein scores. Table SIC lists the background targets as obtained by eluting buffer (instead of citrus protein extract). Note that non-specific targets with low protein scores were obtained by buffer elution. As shown in Fig. 1 (right), the top-ranked citrus targets of P235 and Effector 3 show protein scores far greater than those listed for the non-specific targets in Table sIC. A subset of these targets was further analyzed. The selected P235 targets are SOD (Superoxide Dismutase, from infected citrus), LTP (Lipid Transfer Protein, from healthy citrus), Aspartyl Endopeptidase, (AP) and Glycosyl Hydrolase family 17, GH17 (from both healthy and infected citrus) whereas the Effector 3 targets are: KTI (Kunitz Trypsin Inhibitor) and Aldehyde Dehydrogenase (ALDH) from both healthy and infected citrus, Elongation Factor Tu (Ef-Tu) from infected citrus, lectin, and 21 kDa seed protein-like (a functional homolog of KTI) from healthy citrus. As indicated, all the target proteins listed in Fig. 1(right) are involved in citrus innate immunity. Although, it allows identification of both extracellular and intracellular targets of *CLas* effectors from infected and healthy citrus, our method in Fig. 1 is likely to miss the citrus targets that are expressed at a low level. Most importantly, our *in vitro* method does not prove that the targets listed in Fig. 1 (right) also interact with the *CLas* effectors *in planta*.

**Fig. 1.**
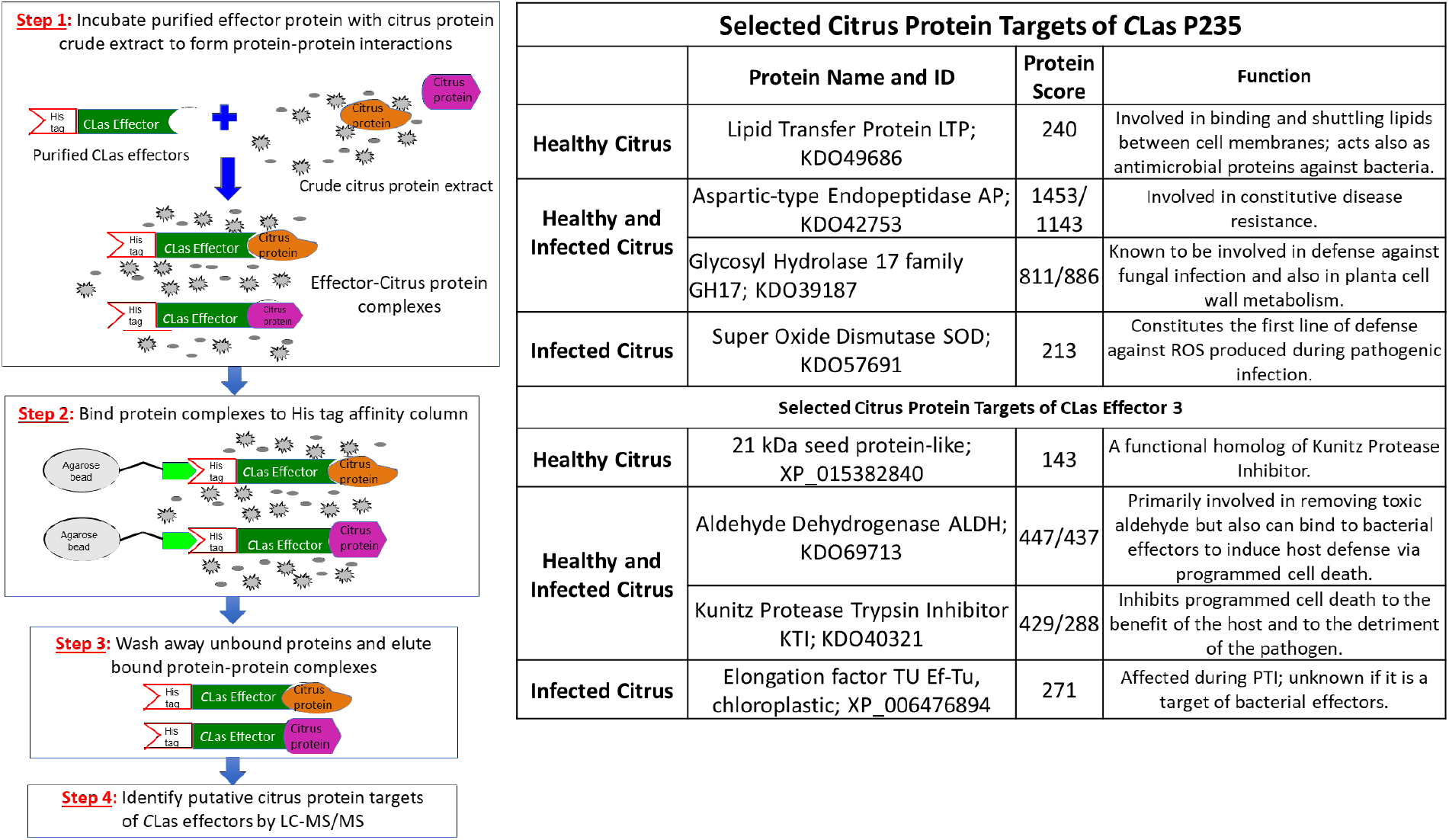
Identification of citrus target proteins of the *CLas* effectors. **(Left)** Outlines of the experimental steps. (Step 1) *CLas* effectors were overexpressed in *E. coli* with His_6_ tag. Purified His_6_-tagged effectors (colored green) were incubated with protein extracts from healthy and infected Hamlin citrus. Specific citrus target protein (colored orange and magenta) bound to the *CLas* effectors. (Step 2) Effector-target complexes were Talon on Agarose beads and non-target citrus proteins were washed away. (Step 3) The specific effector-target complexes were eluted. (Step 4) The citrus target proteins from the eluted complexes were identified by LC-MS/MS. **(Right)** Selected citrus protein targets of the *CLas* effectors, P235 and Effector 3. The citrus targets were chosen on the basis of their high protein scores. The target proteins were selected both from healthy and *CLas*-infected protein extracts. GenBank sequence IDs and putative functions (based upon the literature data) of the citrus targets are listed.

### *In planta* validation of the citrus protein targets of P235 and Effector 3

*In planta* validation is based on a split triple green fluorescent protein (GFP) assay ^56,57^, which has been successfully applied to monitor protein-protein interactions in yeast, human, and plant. The assay relies on the principle that specially enhanced 11 stranded GFP can be split into GFP1-9, GFP10, and GFP11 with none of the three split components showing fluorescence. However, fluorescence is recovered when GFP1-9, GFP10, and GFP11 are re-assembled. We constructed stable transgenic tobacco lines that overexpress GFP1-9 as a detector of *in planta* protein-protein interactions. Two agrobacterium constructs, i.e., one overexpressing P235 (or Effector 3) with a GFP11 tag and the other a putative citrus target with a GFP10 tag, were infiltrated in the GFP1-9 transgenic tobacco. As shown in the experimental design of Fig. 2A, we expect to observe (i) green fluorescence in the presence of a target-effector interaction and (ii) no fluorescence in the absence. In our assay, for negative controls (see Fig. 2B), we confirmed the lack of interaction between Effector 3 and the targets for P235 (and the lack of interaction between P235 and the targets for Effector 3). Agrobacterium carrying enhanced GFP was used as a positive control. Fig. 2C (top) shows the results of the split GFP assay monitoring the interaction of P235. Note the presence of fluorescence at the infiltrated leaf sites for SOD, LTP, AP, and GH17, which were identified as putative targets of P235 from our *in vitro* protein assay as described Fig. 2A. The pattern of fluorescence is comparable to the infiltration of agrobacterium carrying enhanced GFP. Thus, the split GFP assay shows specific *in planta* interactions between *CLas* P235 and citrus proteins (SOD, LTP, AP, and GH17). Fig. 2D (bottom) shows the results of the split-GFP assay monitoring the interaction of Effector 3. The presence fluorescence at the filtrated sites indicates specific *in planta* interactions between Effector 3 and (KTI, ALDH2, lectin, and Ef-Tu) that were identified by the *in vitro* protein assay. Triple split GFP assay provides the following advantages ^58,59^ over other commonly used assays such as yeast-two hybrid system for monitoring protein-protein interaction: (i) it can be readily adapted to *in planta* systems; (ii) it limits false positives and negatives; (iii) small GFP10 and GFP11 tags retain native effector-target interactions; (iv) positive and negative controls can easily be incorporated for *in planta* measurement to improve the fidelity of the assay.

**Fig. 2.**
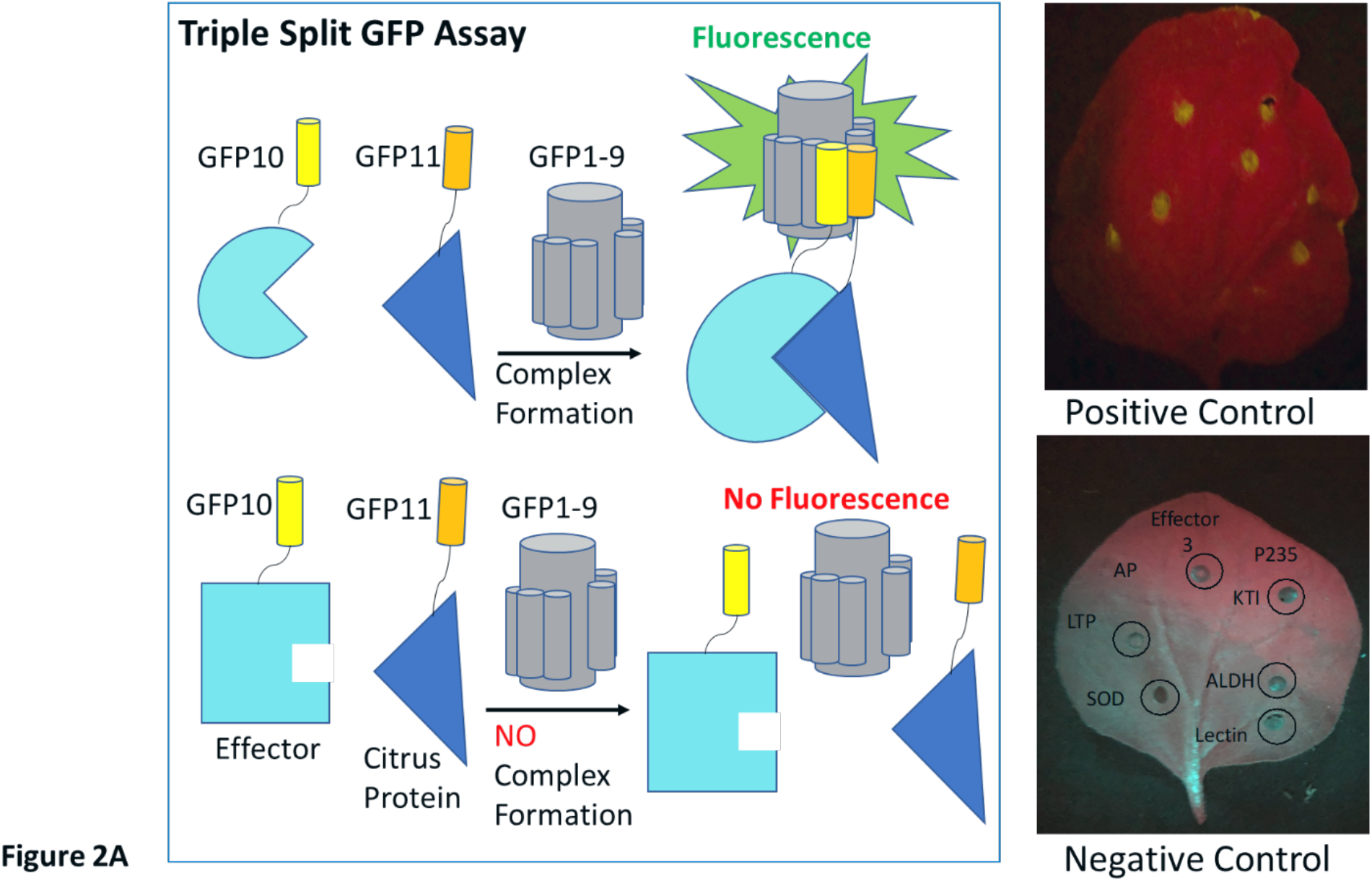
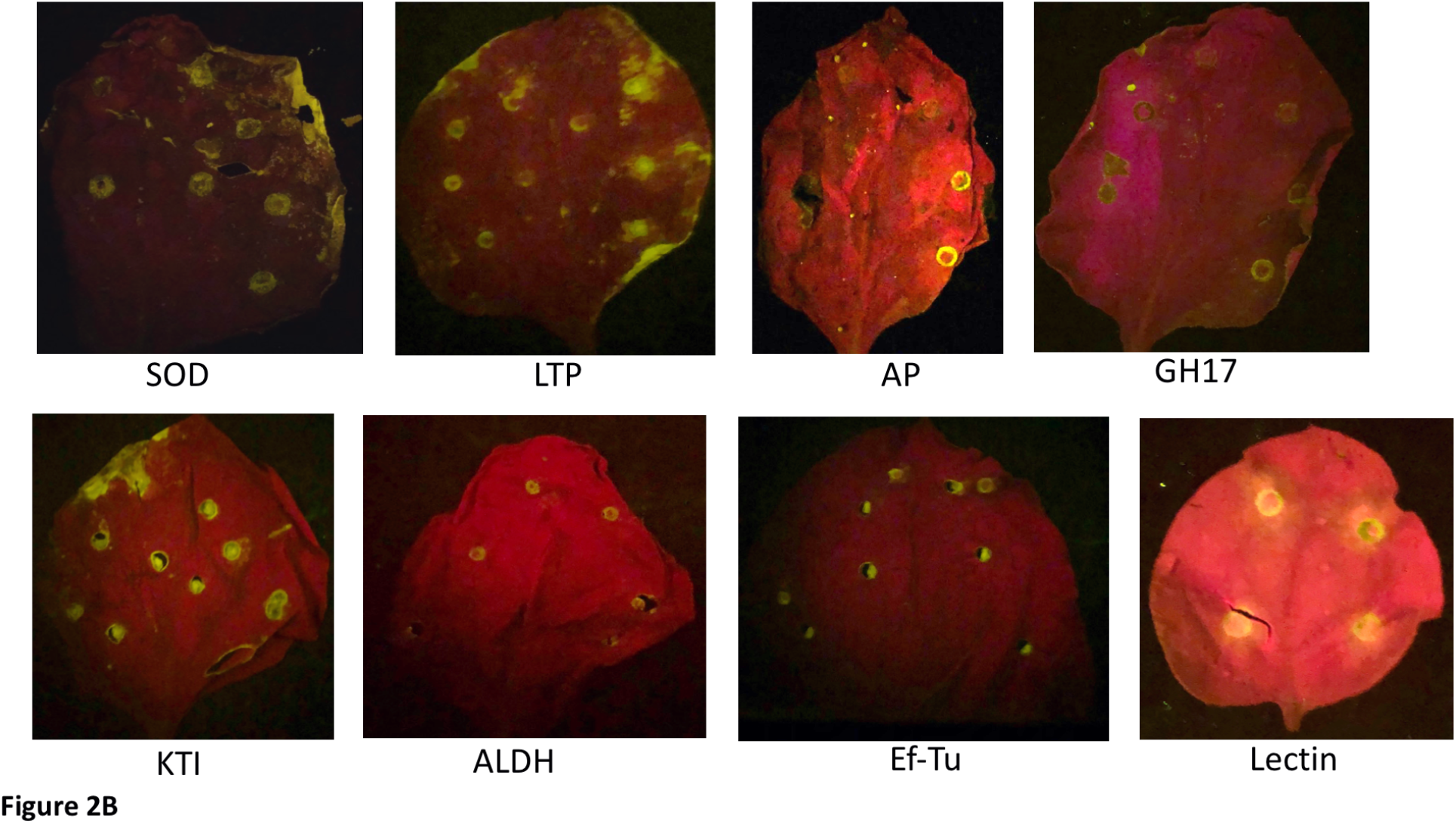
*In planta* validation of the selected citrus protein targets of P235 and Effector 3 by triple split GFP assay. **(A- left)** A schematic representation of the principle of the triple split GFP assay. Green fluorescence is observed when the Effector (cyan) linked to GFP10 (yellow) interacts with the target (blue) linked to GFP11 (orange) and the effector-target complex complements with GFP1-9. There is no fluorescence in the absence of an interaction. **(A- right)** The presence of fluorescence when Agrobacterium carrying enhanced GFP is infiltrated on the leaves of transgenic tobacco expressing GFP1-9 is used as a positive control. Absence of fluorescence when the leaves of transgenic tobacco expressing GFP1-9 co-infiltrated with P235 and the interactors of Effector 3 in one half of the leave (or Effector 3 and the interactors of P235 in the other half of the leave) served as the negative controls. Note the absence of fluorescence. Under the 488 nm excitation filter (blue) and colored glass 520 nm long pass filter, the chlorophyll background appears as red and fluorescence at the site of co-infiltration of the effector and its target appears as greenish yellow spots. (B) Complex formation when P235 is co-infiltrated with SOD, LTP, AP, and GH17 in the leaves of GFP1-9 transgenic tobacco (top panel) and when Effector 3 is co-infiltrated with Effector 3 and KTI, ALDH, Ef-Tu, and Lectin (bottom panel).

### The two *CLas* effectors inhibit the functions of their specific citrus targets

#### *In vitro* assays

Three targets of *C*Las P235, *i.e.*, Fe-SOD, AP, and GH17, that are validated by *in planta* split GFP assay, are citrus PR or defense proteins with enzymatic activities. As described in the “Methods”, the citrus target proteins were expressed in *E. coli*, extracted from the inclusion body, and purified by affinity purification schemes. After purification the proteins are re-folded. Therefore, before conducting *in vitro* inhibition assays, it was necessary to determine the enzymatic activities of the recombinant enzymes to confirm that they retained the native fold and function. We then determined the inhibitory activity of P235 on them by measuring IC50 (the concentration required for 50% reduction in enzymatic activity). Fe-SOD, unique to plants, prevents damage caused by the ROS burst upon pathogen infection^60^. While it facilitates direct killing of the pathogen and induction of plant defense genes, excessive ROS is damaging to the plant. Fe-SOD mainly produced in the cytosol and chloroplast converts oxygen radical to molecular oxygen and hydrogen peroxide. The latter, also potentially phytotoxic, is subsequently converted by plant catalase into molecular oxygen and water. Fe-SOD is also involved in regulating ROS signaling leading to the induction of defense genes ^61^. As shown in Table I, P235 inhibits the activity of the citrus Fe-SOD. Citrus AP belongs to the A1 family of atypical aspartate proteases, primarily located in apoplast and chloroplast. It has been shown that an atypical aspartate protease, expressed in the apoplast, confers constitutive disease resistance 1 (CDR1) in *Arabidopsis* probably by producing a peptide ligand through cleavage and subsequent induction of SA signaling and expression of PR genes ^62,63^. The results of enzyme assay show that P235 inhibits the activity of the citrus AP. GH17, a citrus (β1-3) glucanase, is another direct interactor of P235. Typically, GH17 glucanases are known to provide disease resistance against fungi by hydrolyzing fungal chitins ^64^. But GH17 also has a role in immune defense in general in that it regulates the formation of callose (a β 1-3 glucan polysaccharide), which is an essential component of papillae, an ultrastructure formed at the site of pathogen penetration. Apart from callose, the papillae also contain ROS and antimicrobial peptide thionin and thus provide the first line of defense against pathogen invasion. In papillae-mediated immunity, callose may be involved in two different mechanisms of plant defense against pathogens. *First*, callose deposition in papillae may block pathogen spread. *Second*, hydrolyzed products of callose by GH17 may be ligands for plant PRRs and may induce SA signaling leading to immune defense. Thus, GH17-mediated hydrolysis of callose may either support pathogen spread or induce SA signaling. Since according to the data in Table I, it inhibits the glucanase activity of the citrus GH17, P235 of pathogenic *C*L*as* may suppress citrus immune defense by blocking SA signaling. The citrus LTP is the non-enzyme direct interactor of P235. Plant LTPs possess (i) lipid binding property, which is critical to lipid homeostasis and membrane dynamics and (ii) bactericidal activity as a component of immune defense ^65–68^. Table I shows that P235 can block both lipid-binding and antimicrobial activities. Table I shows inhibitory activities of Effector 3 on two citrus target proteins: ALDH, which converts aldehydes into carboxylic acid using NADPH/NADH as a co-factor ^69^ and KTI, which inhibits protease activity of PCD-inducing trypsin^70^ .

**Table I.** The citrus AP, SOD, GH17, and ALDH were overexpressed in and purified from *E. coli* and enzymatic assays were performed on them following the protocols described in Experimental Procedures. K_cat_ and K_m_ are given for the citrus AP, SOD, GH17, and ALDH. The IC50 (50% reduction in catalytic activity) by the effector is provided as the ratio of the substrate concentration. ^a^Lipid binding assay was performed for LTP to demonstrate the inhibitory effect of P235. ^b^Bactericidal effect of LTP was monitored using two *E. c*oli strains: BL21 and ATCC25922; the corresponding MIC values are in 1M. Addition of P235 at a concentration same as the MIC completely blocks the bactericidal effect of LTP on the two *E. coli* strains. ^c^Effector 3 is added in the Trypsin inhibition assay by KTI to determine the IC50.

| | Citrus targets | $K_{cat}$<br>(s<sup>-1</sup>) | $K_m$<br>(mM) | IC50 | Substrate /ligand concentration (mM) | Target concentration (mM) | MIC (μM)<sup>b</sup><br>(-P235) BL21 | MIC (μM)<sup>b</sup><br>(-P235) ATCC | MIC (μM)<sup>b</sup><br>(+P235) BL21 | MIC (μM)<sup>b</sup><br>(+P235) ATCC |
| --- | --- | --- | --- | --- | --- | --- | --- | --- | --- | --- |
| P235 | AP | 277.2 | 315.8 | 6.30 | 10 | 20 |  |  |  |  |
|  | SOD | 12.45 | 42.3 | 4.37 | 2 | 4 |  |  |  |  |
|  | GH17 | 0.003 | 0.042 | 1.95 | 1 | 25 |  |  |  |  |
|  | LTP |  |  | 1.43<sup>a</sup> | 8 | 20 | 12.5 | 15 | growth | growth |
| Effector 3 | KTI |  |  | 5.60<sup>c</sup> | 4 | 10 |  |  |  |  |
|  | ALDH | 0.094 | 2.131 | 0.86 | 2 | 10 |  |  |  |  |

### *In planta* assays

*In planta* assays for monitoring ROS production, bacterial clearance, and PCD induction were performed in tobacco to examine the inhibitory effect P235 and Effector 3 on their citrus target proteins. Paraquat (PQ) was used for inducing the production of reactive oxygen species (ROS) in tobacco. The ROS level was monitored using a ROS/Superoxide detection assay ^71^. In this experiment, the ROS level induced by Agrobacterium carrying an empty vector (i.e., no gene) plus PQ was normalized to 100%. Note that, infiltration of Agrobacterium carrying citrus SOD reduced the ROS level significantly below 100%. However, as shown in Fig. 3A (left), simultaneous addition of Agrobacteria carrying P235 and citrus SOD showed the elevation in the ROS level proving *in planta* inhibition of citrus SOD by P235. *In planta* bactericidal activity of citrus LTP was monitored by qPCR that showed the reduction of bacterial load in tobacco infected with *Pseudomonas syringae* pv. DC3000. As shown in Fig. 3A (right), Agrobacterium carrying citrus LTP (0.4X10^8^ cfu/ml) reduced the bacterial load to 37%. Increasing Agrobacterium carrying citrus LTP by 10 times (i.e., 0.4X10^9^ cfu/ml) led to the 75% reduction in the bacterial load. The addition of Agrobacterium carrying P235 (0.4X10^8^ cfu/ml) or 10 times of that increased the bacterial load. This proves that P235 is able to block *in planta* the bactericidal activity of the citrus LTP. Fig. S3A shows the Ct values of a *Pst* gene gene (a measure of bacterial load) at different P235 concentrations. *In planta* studies were conducted in tobacco to examine the effect of (Effector 3 – Lectin/Ef-Tu) interactions. As shown in Fig. 3B (left), infiltration of Agrobacterium carrying Effector 3 induced ROS at a high level (85%). The ROS level due to Agrobacterium carrying an empty vector plus Paraquat was set to 100%. Infiltration of Agrobacterium carrying citrus Lectin or Ef-Tu had negligible effect of the ROS level. Co-infiltration of Effector 3 plus lectin or EF-Tu had very little effect of reducing the ROS level induced by Effector 3 alone. However, combination of lectin and Ef-Tu was able to reduce the ROS level induced by Effector 3. In this regard, it is important to note that some bacteria, such as *P. gingivalis*, *M. tuberculosis*, *H. pylori*, and *B. anthracis*, utilize ROS to support their growth and to establish infection ^72^ whereas plant lectin and Ef-Tu tend to inhibit ROS production or ROS-mediated signaling ^73^. It appears that pathogenic *C*Las may use Effector 3 to maintain ROS level that is beneficial to pathogen growth and infection by inhibiting the ROS-inhibitory actions of citrus lectins and Ef-Tu. Paraquat was also used to induce PCD via ROS in tobacco. PCD was monitored by electrolyte leakage ^74^, which was set to 100% as induced by Agrobacterium carrying an empty vector plus PQ. Infiltration of the Agrobacterium carrying Effector 3 induced ∼50% electrolyte leakage, which, as shown in Fig. 3B (Right), was reduced upon infiltration of Agrobacterium carrying citrus KTI. The co-infiltration of Agrobacteria carrying citrus KTI and Effector 3 elevated PCD thereby confirming that Effector 3 is an inhibitor of the citrus KTI.

**Fig. 3.**
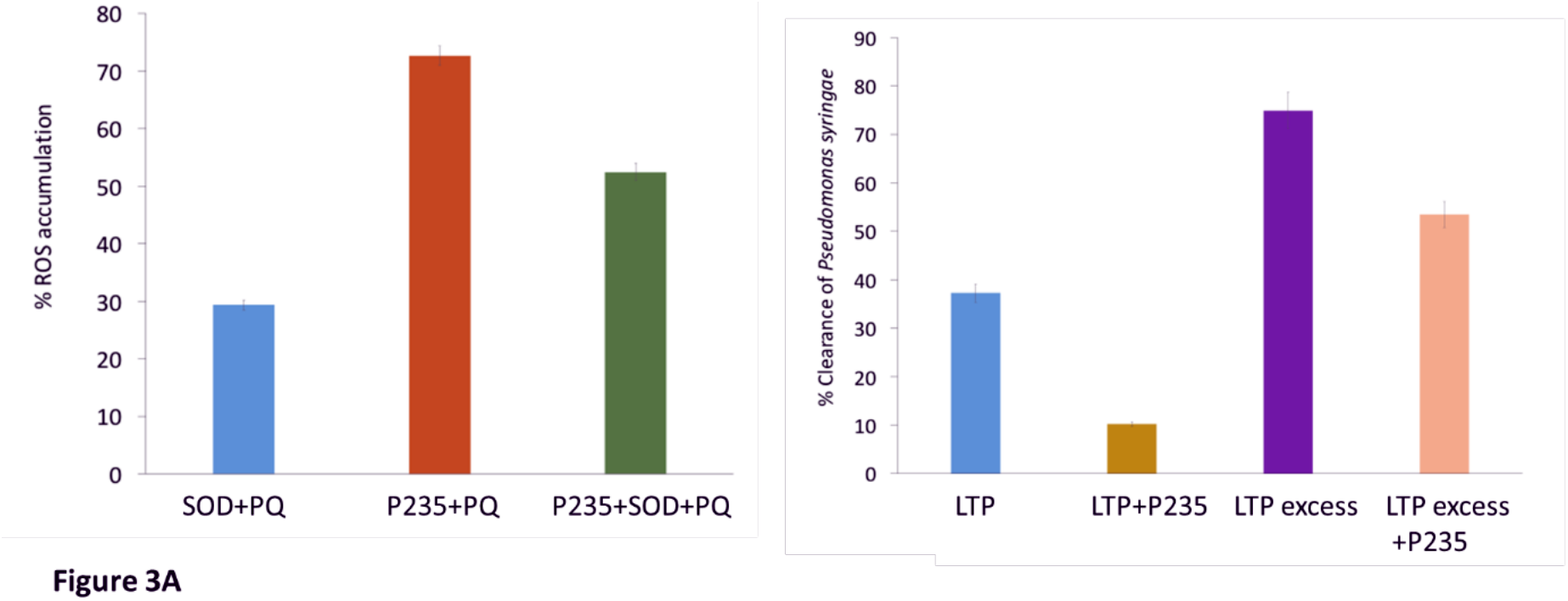
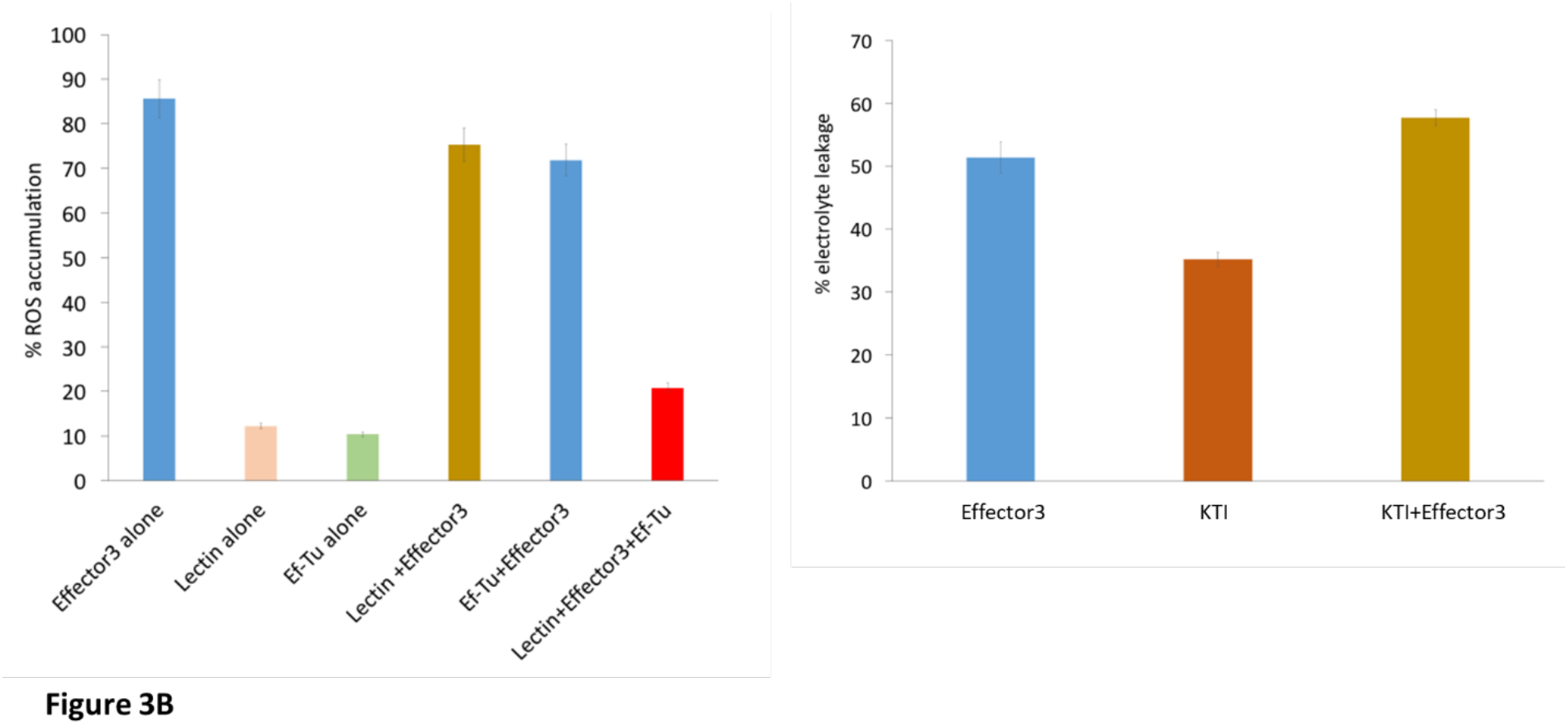
*In vitro* and *in planta* assays to demonstrate the inhibitory activity of the *CLas* effectors on their citrus protein targets. First, enzymatic, binding, or bactericidal assays are performed to determine the appropriate functional properties of the citrus target proteins and subsequently, the same assays are conducted to demonstrate the inhibitory activities of the *CLas* effectors on their citrus targets. **(A- left)** The inhibitory effect of P235 on SOD is monitored in tobacco by fluorescence microscopy. **(A- right)** Percentage reduction relative to the initial cfu of *E. coli* ATCC25922 by LTP alone and LTP plus P235. **(B- left)** The ability of Effector 3 to induce the ROS release in tobacco. The addition of lectin or Ef-Tu is not sufficient enough to suppress the ROS release by Effector 3. However, the combination of lectin and Ef-Tu significantly reduces the ROS release by Effector 3. **(B- right)** The electrolyte leakage due to ROS-produced by PQ is set 100%. Relative electrolyte leakage due to infiltration of Effector 3, KTI, and Effector 3+KTI in tobacco.

### To predict and validate the molecular mechanism of effector-target inhibitory interactions

We performed all-atom molecular dynamics (MD) simulations ^75^ to predict the interactions that stabilize the (inhibitory *CLas* effector-citrus protein target) complexes. Initially, we focused on the citrus LTP vis-à-vis its bactericidal effect. As described in Methods Section, we first obtained an optimized homology-based model of the citrus LTP as shown in Fig. s1C of the supplementary material. Then, we performed MD simulations in (water: lipid) bilayer for 10 µs. As described in the Fig. S3A of the supplementary material, MD simulations revealed that the LTP helices h2, h3, h4 and the C-terminal loop were involved in interaction with the lipid bilayer defining membrane attachment, which is the first step in the bactericidal activity. For the LTP membrane attachment, the interactions of the positively charge arginine residues R21, R32, R39, R44, R71 and R89 (shown in blue in Fig. 4B) with the negatively charged lipid polar heads appear to be extremely critical. In order to study the interaction of P235 with LTP, we docked the homology based P235 model to the optimized LTP model. We then performed MD simulations of the LTP-P235 complex in aqueous environment for 6 µS in order to determine which mode of P235 binding may block the LTP attachment to the lipid bilayer as discussed in Fig. S3A and S3B. One mode of P235 (magenta) interaction, shown in Fig. 4A (left), involves the LTP (cyan) helices h2, h3, and h4 and the C-terminal loop resulting in partial blocking of the B1 LTP site by P235. The prominent pair-wise contacts between P235 (magenta) and LTP (cyan) are predicted from MD simulations using the method described in Fig. S4. They are: S23-R44, P27-R44, R37-F92, F123-R56. Another mode of P235 binding as shown in Fig. 4A (right) involves the LTP helices h1, h2, and h3 with pairwise contacts: I107-R39, R110-A37, R110-Q4, F111-G36. In both binding modes, the LTP attachment to the bacterial membrane is partially blocked. Based upon the two modes of interactions, we designed two LTP mimics shown in Fig. 4B, i.e., Mimic 1 containing h2, h3, h4, and the C-terminal loop and Mimic 2 containing h1, h2, and h3. We also introduced amino acid substitutions, i.e., R44F, R56F, and F92E in Mimic 1 and Q4E, G36F, A37E, R39E in Mimic 2. These amino acid substitutions are predicted to increase the strength of pairwise interactions between LTP and P235 as listed above. While both the Mimics are predicted to partially block the inhibitory activity of P235 on bactericidal LTP, Mimic 1 is supposed be a better blocker than Mimic 2. Our predictions are validated by the results of *in planta* tobacco studies shown in Fig. 4B. Here, Mimics 1 and 2 were infiltrated to express at the same and 10 times level of P235. The results show that: (i) both the mimics by themselves show bactericidal effect on *P. syringae pv.* tobaci but smaller than the full-length LTP; and (ii) Mimic 1 is better P235 blocker/bactericidal than Mimic 2. These experimental observations are in full agreement with our predictions. Therefore, we may conclude that interactions shown in Fig. 4B (left) is the most prominent mode of LTP blocking by P235. Fig. s4B shows the Ct values of a *Pst* gene (a measure of bacterial load) due to treatment of Mimics 1 and 2 at different concentrations.

**Fig. 4.**
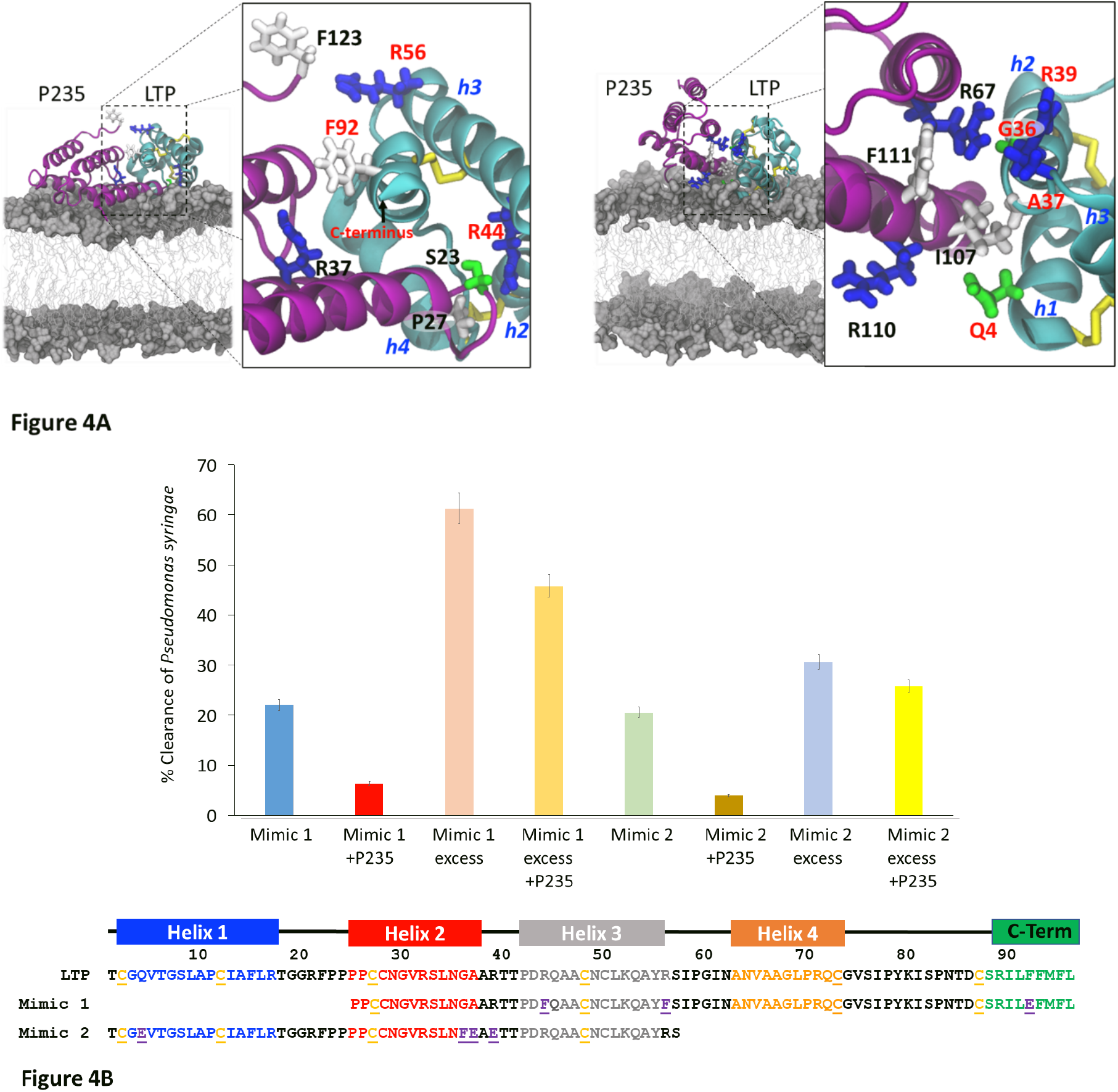
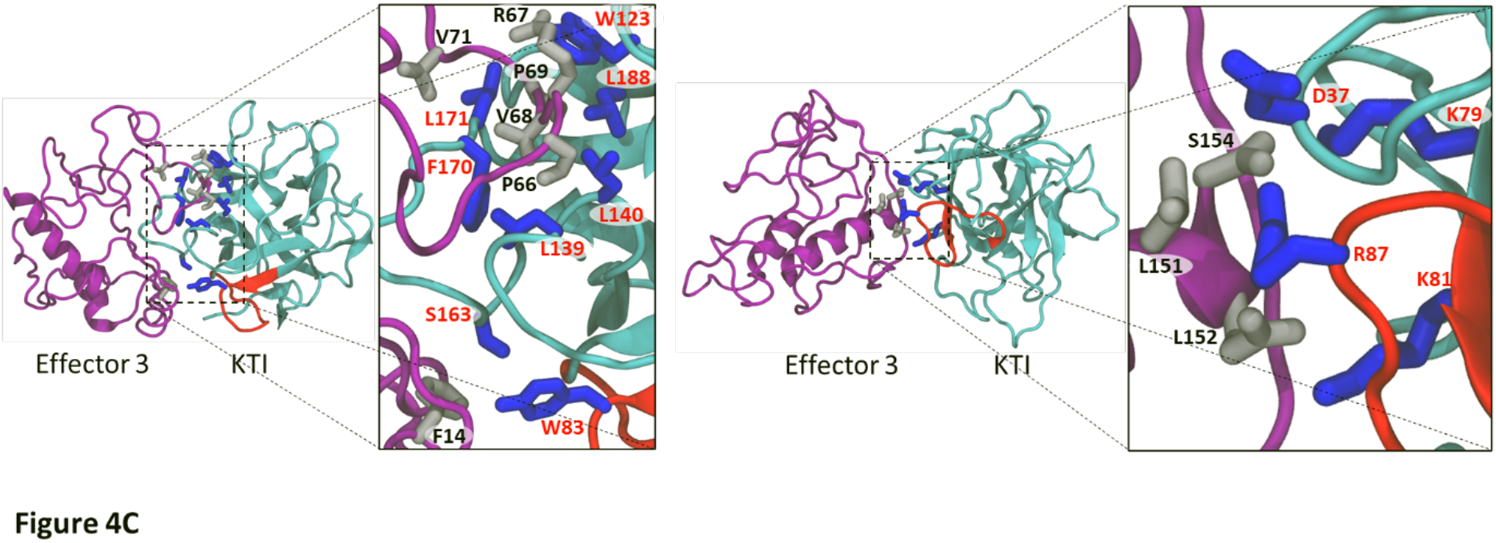
Prediction and validation of (citrus target-*CLas* effector) interaction at the contact interface. Molecular modeling is performed to predict the pairwise interactions based on which mimics are designed to displace the effector from the (citrus target-*CLas* effector) complex. Finally, experiments are performed to determine if the mimic, indeed, displaces the effector from the complex and if so, our prediction of the pairwise interactions are validated. **(A)** Two possible modes of interactions between P235 (magenta ribbon) and LTP (cyan ribbon) which block the membrane attachment of LTP thereby inhibiting the bactericidal activity. The disulfide bridges are shown as yellow sticks. The LTP helices (h2, h3, and h4) and the C-terminal segment are predicted to be involved in one mode of interaction. The LTP helices (h1, h2, and h3) are involved in the other mode of interaction. Important residues in the pairwise contacts are shown: P235 residues are labeled black whereas the LTP residues are labeled red. Basic, acidic, neutral, and acidic residues are respectively as blue, red, green, and grey sticks. **(B)** Amino acid sequences of Mimic 1 and Mimic 2 at the bottom. Bacterial (*P. syringae* pv. tobaci) clearance in tobacco by the two mimics in the presence and absence of P235. Note that an excess of the mimics is needed for significant bacterial clearance. Mimic 1 is a better bactericidal than Mimic 2. P235 is an inhibitor of Mimic 1 or Mimic 2. **(C)** Two models of interactions between Effector 3 (magenta ribbon) and KTI (cyan ribbon with the reactive loop in red). Both the models show interactions with the KTI reactive loop as a prominent mode of inhibition. The predicted pairwise interactions are shown. The residues from Effector 3 are shown as grey sticks and labeled black whereas the residues from KTI are shown as blue sticks and label red.

We constructed two models of (Effector 3: KTI) complex with Effector 3 and Kunitz represented respectively by purple and cyan ribbons. Both the complexes are chosen to block the reactive KTI loop (residues 82-94) as shown in the homology-based model of Fig. S1D of the supplementary material. Blocking of the KTI reactive loop is critical in trypsin protease inhibition. We performed 2 µS MD simulations on these two complexes in aqueous environment. Fig. 4C shows two different models that represent two different ways Effector 3 may block the KTI reactive loop (shown in red). The sampling of the MD trajectories reveals the following dominant pairwise interactions with Cœ-Cœ distance < 4Å as described in Fig. S5. Pairwise interactions in one model in Fig. 4D (left) are: F14-W83, P69-W123, V68-L139, V68-L140, F14-S163, V68-F170, L71-L171, P69-L188 whereas in the other model in Fig. 4D (right) they are: L151-D37, L151-R87, L152-K81, S154-K79. In these pairwise interactions, Effector 3 and KTI are respectively shown respectively as grey and grey and blue ball-and-stick representations. Mutational studies are needed to discriminate the two modes of inhibition of KTI by Effector 3 described in Fig. 4D.

## Discussion

Bacterial effectors are often described as inhibitors of plant innate immune signaling networks mediated by PTI, ETI, and plant SA, JA, and ET hormones. The end products of PTI, ETI, and plant hormone signaling are the immune defense proteins that either clear the invading the pathogen or block the infection. Typically, each immune defense protein is induced at a low level and a single protein, therefore, can neither completely clear the pathogen nor can it block the infection. Interestingly, simultaneous induction of multiple immune defense proteins (albeit at low levels) can lead to effective clearance of the invading bacteria and blocking of infection caused by them. However, multiple effectors from a pathogenic bacterium like *C*Las can suppress multiple signaling steps to support bacterial growth and infection. Here, we report the role of two *C*Las effectors, P235 and Effector 3, in HLB pathogenesis. Each of them may directly target and inhibit more than one citrus innate immune defense proteins belonging to the bactericidal and/or disease-blocking proteins. For example, P235 can inhibit the citrus targets (SOD, AP, GH17, and LTP) whereas Effector 3 can inhibit citrus targets (KTI, ALDH, Lectin, and Ef-Tu). Although, as shown here, a bacterial effector may target several plant proteins, inhibitions of all the targets may not be equally important for bacterial pathogenesis. A direct evaluation of the importance of each (plant protein-bacterial effector) interaction is traditionally obtained by knockout of a specific bacterial effector. Since *C*Las is not culturable, it is not possible to conduct gene knockout experiments. However, the inhibitory activities of a *CLas* effector against different citrus targets reveal qualitatively the relative importance of different inhibitory (*C*Las effector–citrus target) interactions in HLB pathogenesis. For example, as shown in Table I, P235 is a potent inhibitor of LTP because at equimolar concentration it can completely block the the bactericidal activity of LTP. Thus, P235 may play an important role in HLB pathogenesis. Note that, relatively low IC50 values (within 1 to 6) in Table I, argue that the corresponding inhibitory interactions may be relevant in HLB pathogenesis. Fig. 5 schematically summarizes the combined effect of the inhibitory interactions of P235 and Effector 3 on their citrus targets as determined from our *in vitro* and *in planta* studies. The immune stimulatory defenses exerted by the identified citrus targets are marked by green lines whereas the inhibition of these targets by the two effectors P235 and Effector 3 of pathogenic *CLas* are marked by red lines. Note that SOD reduces the level of ROS whereas Ef-Tu, Lectin, and ALDH tend to control the toxic damage due to ROS. GH17 and AP provide immune defense via SA-signaling, which may involve ROS production whereas KTI may prevent premature ROS-induced PCD and P235 may block *CLas* clearance by LTP. Thus, P235 and Effector 3, target and interact with the ROS, PCD, and bactericidal pathways in a way that adversely affect citrus innate immune defense and in turn, facilitate HLB pathogenesis.

**Fig. 5.**
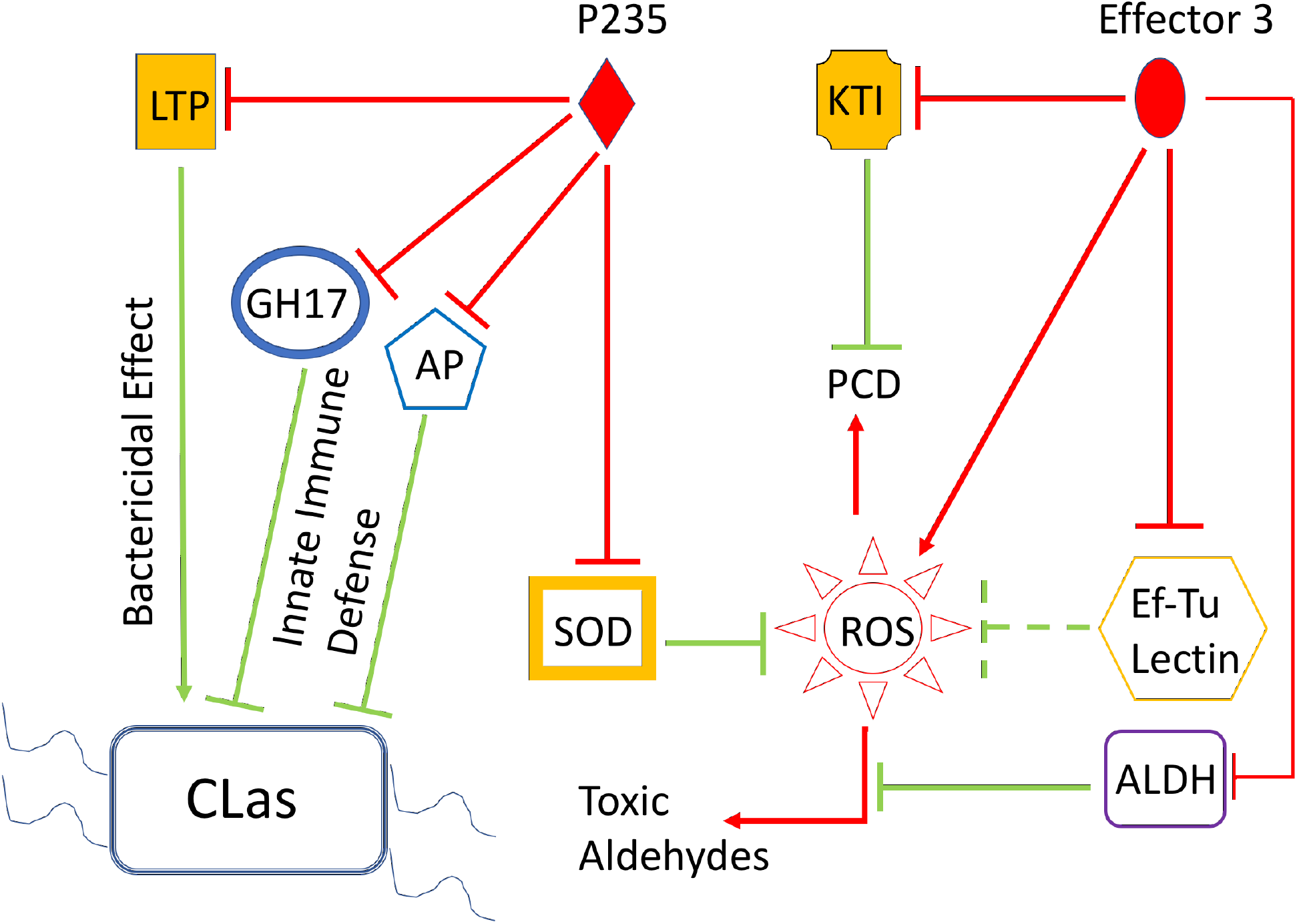
Combined effect of the *CLas* effectors, P235 and Effector 3, on the citrus innate immune due to the target proteins. The P235 targets shown here are: SOD, LTP, KP, and GH17 whereas the Effector 3 targets are: ALDH, Lectin, KTI, and Ef-Tu. All these citrus targets participate in innate immune defense (shown as green arrows) during bacterial infection. For example, SOD controls the level of ROS such that the beneficial effects of ROS-induced immune defense can be harnessed without the level of ROS level exceeding a critical threshold over which there may cause host damage. EF-Tu and ALDH also limit the level of ROS. GH17 and KP offer immune defense against bacterial infection. LTP can directly exert bacterial effect whereas KTI prevents premature PCD, which may help bacterial growth and infection. The effects of pathogenic *CLas* effectors, P235 Effector 3, are shown as red line arrows (promoting a process and red line blockers (as inhibiting a process). The combined effects of P235 and Effector 3: elevation of ROS level, premature PCD, and inhibition of *CLas* clearance.

We analyzed the detailed interactions at the contact interfaces of the (P235-LTP) and (Effector 3-KTI) complexes. Molecular modeling and mutational analysis revealed the predominant mechanism of LTP inhibition by P235. We were able to design Mimic 1 (derived from LTP with specific amino acid substitutions) that showed intrinsic bactericidal activity and exhibited P235 inhibitory activity. The Mimic 1 can be further modified to increase its P235 inhibitory and bactericidal activity. We have also obtained two modes of inhibition in which Effector 3 may block the reactive loop of the citrus KTI. We have not yet completed *in plant*a experiments to determine whether one of the two modes of inhibition or both may be important. Nonetheless, the citrus KTI as the target of inhibition by a *C*Las effector is an interesting observation since such inhibition may cause premature PCD, which may be beneficial to *C*Las in causing infection ^76^.

## Acknowledgments

This work was funded by the NIFA-CDRE grant (Fed Award # 2018-70016-27453 to PI: Jinhe Bai, USDA-ARS; Co-PI: Subaward # 59-6034-8-005) on “Accelerating implementation of HLB hybrids as new commercial cultivars for fresh and processed citrus (PI:) through a collaboration between Gupta (New Mexico Consortium) and Stover (USDA-ARS) labs. We thank Dr. Steven Buelow (Director, New Mexico Consortium) for his support and access to the resources and facilities of the Biolab. We are grateful to Dr. Geoffrey Waldo for generous gifts of plasmids and enhanced GFP used for the split GFP assay.

## Author contributions

G.G. contributed to project planning, experimental work, data analysis, and writing of the manuscript. S.B. performed the majority of *in vitro* and *in planta* studies, data analysis and writing of the manuscript. L.H. performed the MD simulations and analyses. H.N. assisted in the split GFP assays. R.R. performed expression of some recombinant proteins. J.V. performed some MIC assays. E.S. supervised the work of G.M. and Q.S. on citrus infection by *CLas* and protein extraction from healthy and infected citrus and the work of G.H. and S.Z. toward generating transgenic tobacco expressing GFP1-9.

## Supplemental Information

Supplemental Information is attached below.

## SUPPLEMENTARY INFORMATION

**Fig. S1.**
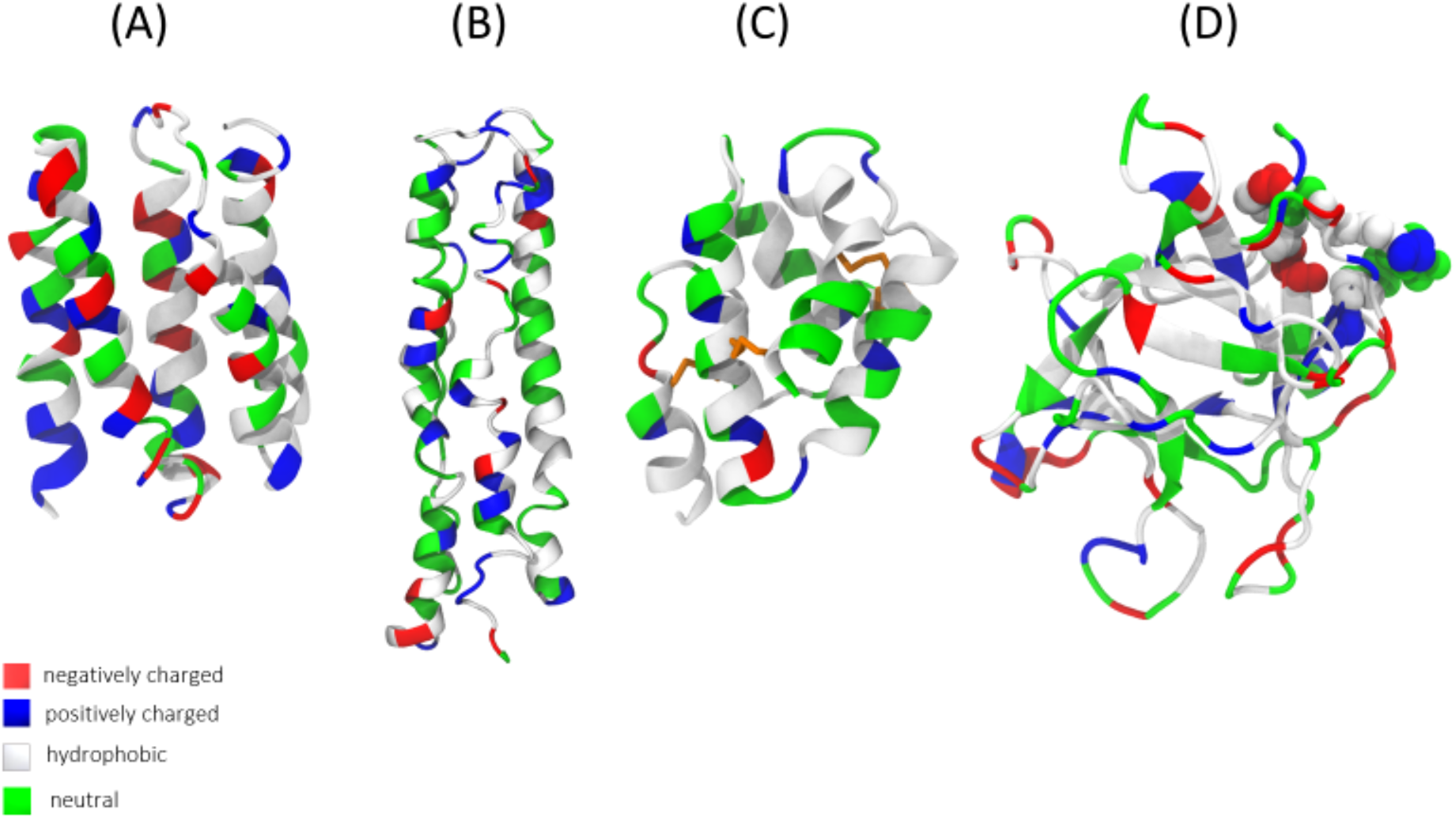
Homology based and energy-minimized models of **(A)** P235, **(B)** Effector 3, **(C)** LTP, and **(D)** KTI, the reactive loop of which is shown as space-filling representation.

**Fig. S2.**
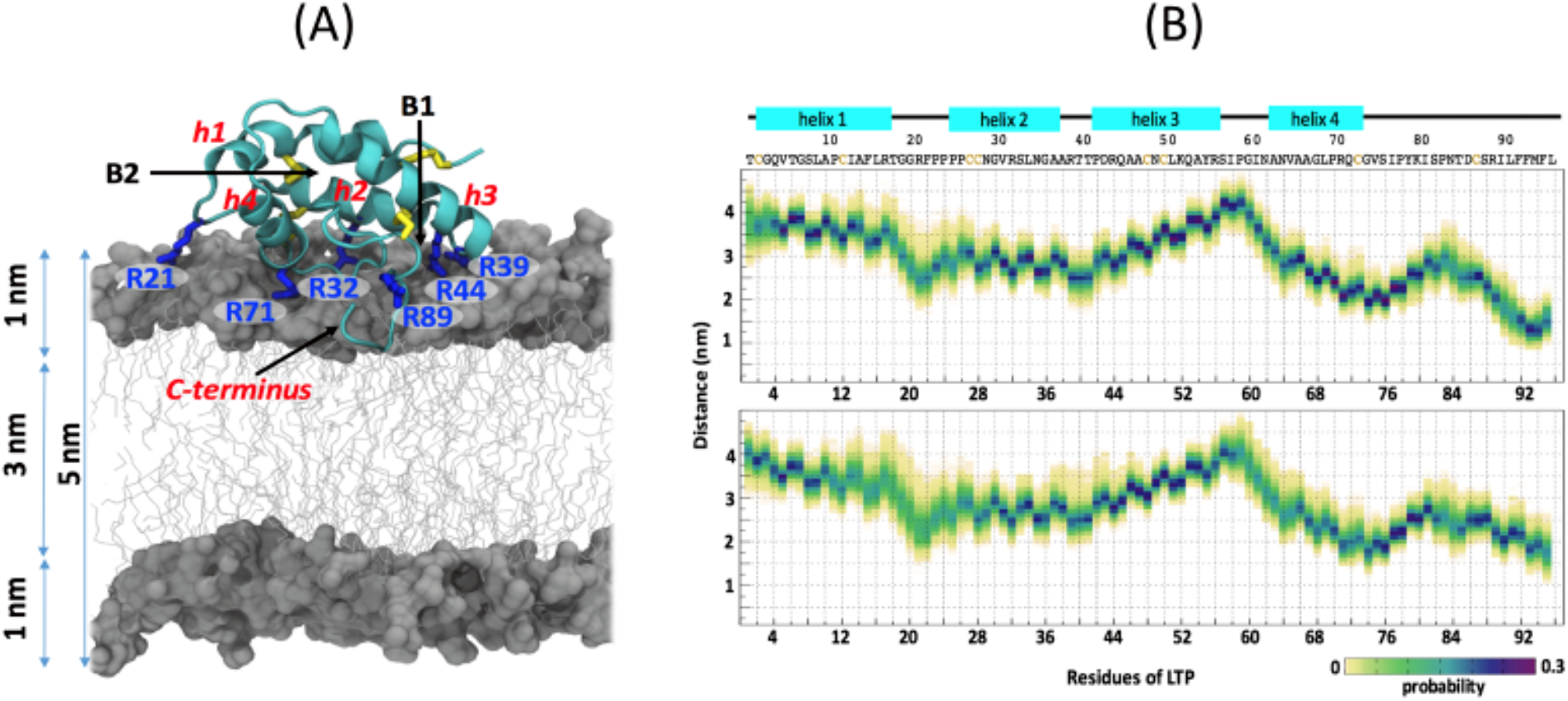
Western blot analysis of the citrus targets for **(A)** P235 and **(B)** Effector 3 by an anti-His_6_ antibody. In (A) Lanes, 1: Marker; 2: Aspartyl Protease; 3: Glycosyl Hydrolase 4: Superoxide Dismutase; 5: Lipid Transfer Protein. In (B) Lanes, 1: Marker; 2: PSII subunit protein; 3: Aldehyde dehydrogenase; 4: Kunitz trypsin inhibitor; 5: Lectin like protein.

**Fig. S3.**
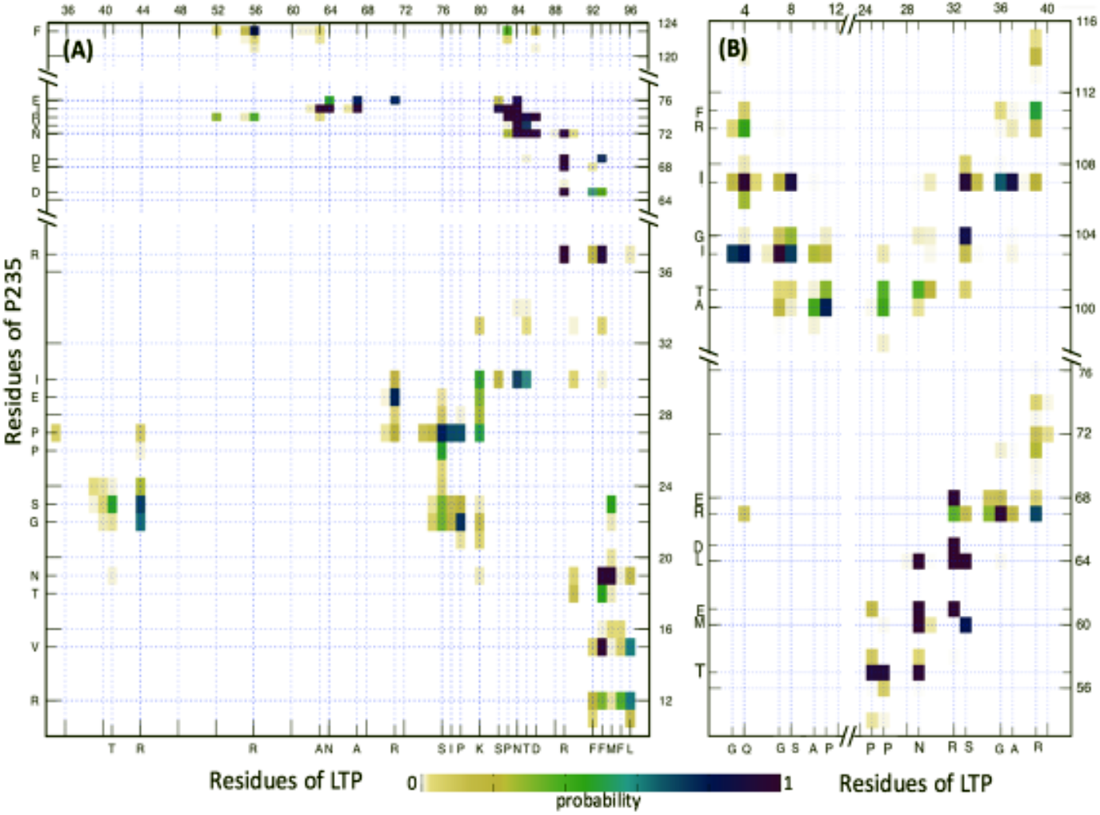
**(A)** Schematic representation predicted models of interaction between LTP and lipid bilayer. Grey lines and surfaces represent lipid acyl chains and head groups, respectively. LTP is shown as a cyan ribbon. LTP-bilayer interaction involves LTP helices h2, h3, h4 with C-terminus segment inserting into the bilayer. Our MD simulations suggest that positively charged residues R21, R32, R39, R44, R71 and R89 (blue sticks) are critical for the interaction with the lipid bilayer. One of LTP lipid entrance sites (B1) is formed by C-terminus and the beginning of h3 with R44 interacting with bilayer membrane. The other LTP lipid entrance site (B2) is formed by C-terminus and loop connecting h3 and h4 and are solvated by water. Other residues at the LTP-lipid interface and water are not shown. Total simulation time was 1-ms. Disulfide bridges for the pairs C2-C50, C12-C27, C28-C73, and C48-C87 are represented with yellow sticks. Amino acid sequence for LTP with residues involved in disulfide bond are underlined: T<u>C</u>GQVTGSLA P<u>C</u>IAFLRTGG RFPPPPCCNG VRSLNGAART TPDRQAA<u>C</u>NC LKQAYRSIPG INANVAAGLP RQ<u>C</u>GVSIPYK ISPNTD<u>C</u>SRI LFFMFL. **(B)** Heatmaps depict residue-specific distributions of the distance between each C_α_ atom and the bilayer center along its normal of LTP-membrane separation. Two independent molecular simulations of LTP-membrane (top and bottom) were conducted with random initial location of the LTP in water. Dashed red line at 2.0 nm indicates the average position of lipid phosphorus atoms. Residues that from disulfide bridge are in yellow (C2-C50, C12-C27, C28-C73, and C48-C87). Both (top and bottom) MD simulations show part of helix 4 and C-terminus of LTP insert deeper into the bilayer, with helix 2 and the beginning of helix 3 interacting with lipid head groups. As shown, one of LTP lipid entrance sites is formed by C-terminus and the beginning of h3 with R44 interact with bilayer membrane. The other LTP lipid entrance site is formed by C-terminus and loop connecting h3 and h4. The C-terminal residues are either in the disordered loop or helical conformation. The total simulation time was 10 ms per system.

**Fig. S4.**
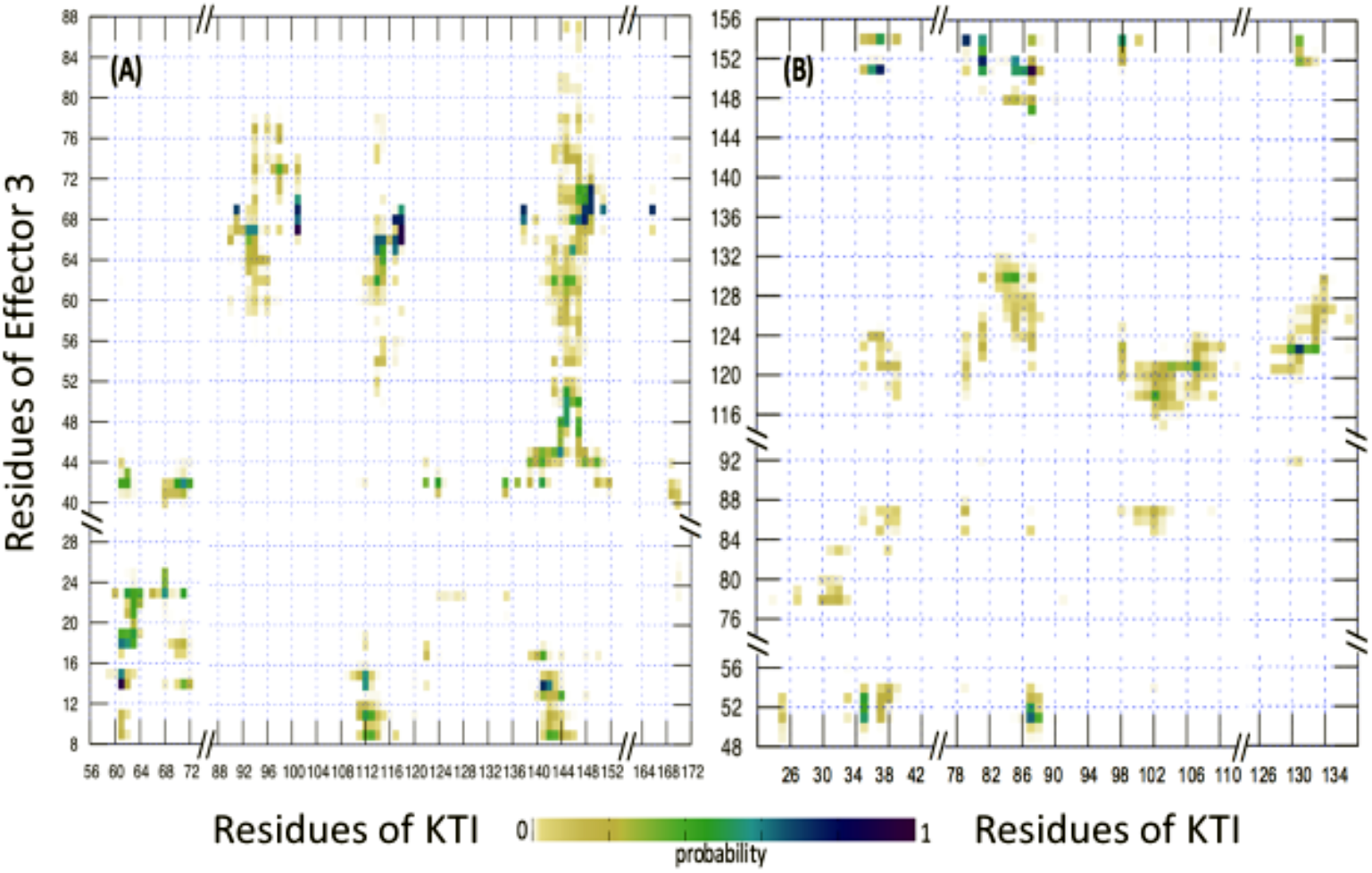
Heatmaps depict pairwise residues interaction between C_α_ atoms of P235 and LTP with C_α_-C_α_ interaction distance ≤ 4.0 Å. MD simulation of P235-LTP were conducted in the presence of lipid bilayer. P235 either interacts with **(A)** helix 2, 3, 4 and the C-terminal segment with or **(B)** helix 1, 2, and 3 of LTP.

**Fig. S5.** Heatmaps depict pairwise interactions between C_α_ atoms of Effector 3 and KTI residues with C_α_-C_α_ interaction distance ≤ 4.0 Å. Active loop of KTI is comprised of residues 82 to 94. Signaling sequence of KTI (residues 1-22) was not included in the simulation model. MD simulations of the (Effector 3-KTI) complex were conducted in the presence of water. Two modes of interaction were predicted: **(A)** minimal interactions of Effector 3 with the KTI active loop residues and **(B)** direct interactions of Effector 3 with the KTI active loop residues, R87 of KTI.

**Table SI.** An expanded list of putative citrus targets of **(A)** CLas-P235, **(B)** CLas-Effector 3, and **(C)** control buffer obtained following the method described in Fig. 1 in the main text.

| Healthy (Unique) | Protein Name | Score | Amino Acid Length |
| --- | --- | --- | --- |
| KDO49686 | lipid-transfer proteins | 240 | 126 |
| XP_006473709 | thioredoxin | 208 | 238 |
| KDO37517 | metal ion binding | 135 | 139 |
| KDO44469 | stromal 70 kDa heat shock-related protein | 113 | 714 |
| KDO73834 | groES | 113 | 253 |
| KDO66479 | lactoylglutathione lyase | 64 | 362 |
| KDO67640 | HSPA1-2_6-8 | 197 | 515 |
| KDO41601 | beta-acetyl hexosaminidase | 251 | 596 |
| **Common Ones (Healthy/Infected)** |  |  |  |
| KDO49891 | Fructose-bisphosphate aldolase | 1549/1196 | 398 |
| KDO69145 | ATP binding | 1045/1171 | 465 |
| KDO42753 | **aspartic-type endopeptidase activity** | **1453/1143** | 423 |
| KDO39187 | **glycosyl hydrolase 17 family** | **811/886** | 348 |
| KDO61594 | Phosphoglycerate kinase | 823/883 | 474 |
| KDO61597 | Phosphoglycerate kinase | 221/307 | 401 |
| XP_006471877 | glutathione synthetase | 362/853 | 573 |
| XP_006495232 | Lectin | 517/694 | 268 |
| KDO61018 | phosphopyruvate hydratase activity | 556/650 | 445 |
| KDO85798 | transketolase activity | 474/582 | 748 |
| KDO85095 | CATALASE | 468/576 | 598 |
| KDO69713 | NAD(P)+-dependent aldehyde dehydrogenase | 589/423 | 456 |
| KDO66162 | carbonic anhydrase | 359/421 | 334 |
| KDO40321 | endopeptidase inhibitor | 226/397 | 211 |
| KDO60185 | chaperonin (HSP60) family | 498/392 | 582 |
| KDO72844 | triose-phosphate isomerase | 297/324 | 173 |
| XP_006488950 | chaperonin (HSP60) family | 151/288 | 608 |
| KDO61040 | Fructose-bisphosphate aldolase | 217/276 | 358 |
| KDO37191 | Fructose-bisphosphate aldolase | 233/274 | 383 |
| KDO85472 | nucleoside diphosphate kinase | 146/269 | 152 |
| KDO78261 | Lactoylglutathione lyase | 218/259 | 291 |
| XP_006488297 | heat shock protein 70 family | 259/242 | 649 |
| XP_006473724 | Cupredoxin | 429/168 | 168 |
| XP_006473831 | GAPDH | 231/229 | 453 |
| **Infected** |  |  |  |
| KDO50492 | peroxisomal (S)-2-hydroxy-acid oxidase | 317 | 370 |
| KDO61080 | Pyridine nucleotide-disulphide oxidoreductase | 213 | 461 |
| KDO57691 | **SOD** | **213** | 242 |
| XP_006480766 | heat shock protein 70 family | 132 | 647 |

Table sIA
| Healthy (unique) | Protein Name | Scores | Amino Acid Length |
| --- | --- | --- | --- |
| KDO80090.1 | Glycine Decarboxylase | 293 | 1049 |
| KDO73087.1 | Heme Dependent Peroxidases | 166 | 265 |
| KDO53866 | flavonoid reductase | 151 | 245 |
| XP_015382840 | 21 kDa seed protein-like | 143 | 159 |
| KDO76988 | RPLO | 134 | 266 |
| XP_006466404 | cysteine desulfurase | 117 | 457 |
| **Common (Healthy/Infected)** |  |  |  |
| XP_006495232.1 | lectin | 635/1774 | 268 |
| KDO69713 | Aldehyde Dehydrogenase 2 | 447/437 | 456 |
| KDO40321 | Kunitz PTI | 429/288 | 211 |
| XP_006475931 | Ascorbate Peroxidase | 293/156 | 1058 |
| XP_006492335 | CP12-2 | 104/181 | 130 |
| **Infected** |  |  |  |
| KDO69145 | ATPase | 778 | 465 |
| KDO42753 | pepsin_A_like_plant protease | 619 | 423 |
| KDO61018 | enolase | 589 | 445 |
| XP_006475931 | Glycine Decarboxylase | 280 | 1058 |
| XP_006476894 | Ef-Tu | 271 | 482 |
| KDO82251 | Alanine Transaminase | 228 | 463 |

Table sIB
| Healthy | Score | Protein Name |
| --- | --- | --- |
| XP_006484037 | 164 | Glyceraldehyde-3-phosphate dehydrogenase |
| XP_006493767 | 103 | Glyceraldehyde-3-phosphate dehydrogenase |
| XP_006474808 | 169 | Fructose-bisphosphate aldolase |
| AFF58923 | 139 | Hsp90 OS |
| XP_006467413 | 138 | Aminomethyltransferase |
| XP_006492777 | 38 | Beta-hexosaminidase |
| XP_006482343 | 38 | 6-phosphogluconate dehydrogenase |
| XP_024957097 | 31 | Carbonic anhydrase |
| XP_006492442 | 24 | Peptidyl-prolyl cis-trans isomerase |
| XP_006488027 | 20 | Lactoylglutathione lyase |
| Infected | Score | Protein Name |
| XP_006477480 | 47 | Ribulose-phosphate 3-epimerase |
| XP_006470328 | 117 | Carbonic anhydrase |
| XP_006487372 | 46 | Fructose-bisphosphate aldolase |
| XP_006487422 | 41 | peptidyl-prolyl-cis-trans isomerase |
| XP_006467420 | 41 | Lactoylglutathionelyase |
| XP_006493789 | 33 | Phosphoglycerate kinase |
| XP_006481103 | 22 | Ferritin |
| XP_006488930 | 17 | Tubulin alpha |

Table sIC

**Table S2.**
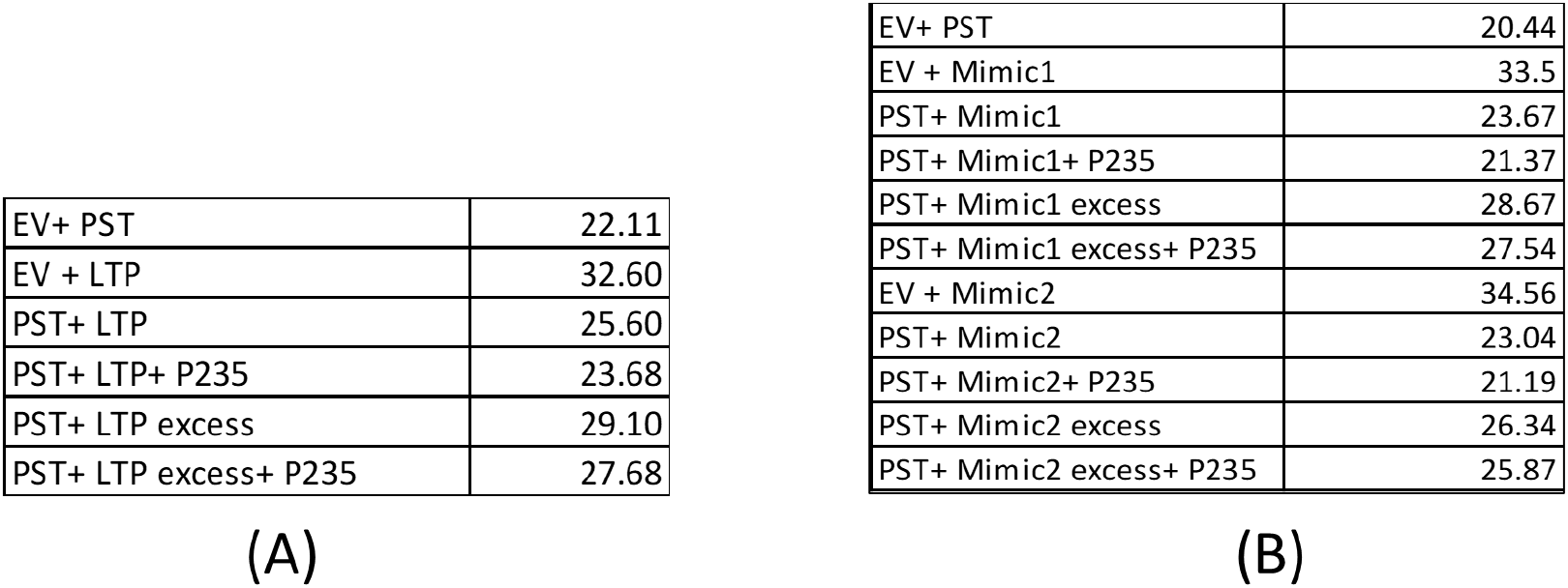
Ct values obtained from qPCR analysis after infiltration of *N.benthamiana* leaves with (A) For LTP +P235 and (B) LTP +P235+ Mimic1 and M2.

Table sIC
|  |  |
| --- | --- |
| EV+ PST | 22.11 |
| EV + LTP | 32.60 |
| PST+ LTP | 25.60 |
| PST+ LTP+ P235 | 23.68 |
| PST+ LTP excess | 29.10 |
| PST+ LTP excess+ P235 | 27.68 |
| (A) |  |
| EV+ PST | 20.44 |
| EV + Mimic1 | 33.5 |
| PST+ Mimic1 | 23.67 |
| PST+ Mimic1+ P235 | 21.37 |
| PST+ Mimic1 excess | 28.67 |
| PST+ Mimic1 excess+ P235 | 27.54 |
| EV + Mimic2 | 34.56 |
| PST+ Mimic2 | 23.04 |
| PST+ Mimic2+ P235 | 21.19 |
| PST+ Mimic2 excess | 26.34 |
| PST+ Mimic2 excess+ P235 | 25.87 |

